# Genetic structure analysis on leopard cat reveals the effects of habitat fragmentation on subpopulation differentiation in Beijing

**DOI:** 10.1101/2022.02.28.482220

**Authors:** Yang Teng, Jing Yang, Longfei Ju, Lixin Gai, Jian Li, Fuli Gao, Weidong Bao

## Abstract

In the face of habitat shrinkage and segregation, the survival of wild cats looks bleak. Interpreting the population genetic structure during habitat fragmentation is critical to planning effective management strategies. To reveal the segregation effects of road construction and human settlement on population genetic structure, we noninvasively analyzed fecal DNA samples from leopard cats (*Prionailurus bengalensis*) from five nature reserves in mountainous areas around Beijing. We focused on molecular microsatellite markers and the mitochondrial control region. A total of 112 individual leopard cats were identified among 601 samples of scat, and generally moderate population genetic diversity was detected. Microsatellite marker–based gene flow (Nm) and STRUCTURE analyses showed no obvious discrepancies among the subpopulations; however, the calculation of genetic distance and construction of a phylogenetic tree of the mitochondrial control region revealed a trend toward differentiation from the other subpopulations in the Songshan subpopulation, which indicates habitat fragmentation effects. Because the isolated Songshan subpopulation may evolve independently, we suggest that its genetic structure be monitored every 5 years to detect any changes and that individuals be introduced as needed to maintain the viability of this subpopulation.

## INTRODUCTION

Variation in population genetic structure can influence the interactions between a species and their environment, and populations with higher genetic diversity are more adaptable to risks brought about by changing environments^[1]^. Moreover, genetic diversity may reflect species’ evolutionary potentiality and supply important information about their current status and conservation strategies^[2]^. When habitat fragmentation occurs, wild animals are threatened by population isolation and genetic loss that may increase the risk for extinction among segregated populations^[3]^. Thus, analyses of genetic structure are becoming an important part of effective conservation^[4]^. Studies on genetic variation may identify subpopulation differentiation, which would help create different management units to maintain the long-term survival of local populations^[5-6]^.

Although small in number, wild feline species are important predators with various body types, diverse food habits, and great adaptability to the surrounding environment. These species play key roles in the natural ecosystem^[7-8]^. As a result of severe disturbance due to human activity, suitable habitats for wild cats are gradually being lost and their prey are also decreasing, which further threatens their survival^[9]^. Most feline species are secretive and highly vigilant, which makes them a very difficult target for field research and conservation^[10-11]^. With the rapid development of noninvasive sampling among wild animals, fecal samples are becoming informative objects in research on wild cats^[12-13]^. Sex ratio determination, individual identification, and relatedness analysis enable accurate evaluation of population genetic diversity and contribute much to the conservation of wild cats^[14-15]^.

The leopard cat (*Prionailurus bengalensis*) is a small wild cat widely distributed in East Asia, South Asia, and Southeast Asia^[16]^. This small cat appears almost everywhere in China except at high altitudes in the Tibetan plateau and in the severe dry lands of the northwest^[17]^. Although it is generally classified as of Least Concern in the International Union for Conservation of Nature (IUCN) Red List of Threatened Species^[18]^, the leopard cat population is decreasing dramatically as a result of habitat loss and illegal hunting^[19]^. Thus, it is listed in the second category of the CITES Appendices to strengthen protection^[20]^. Subspecies of the leopard cat are classified differently in the in the IUCN Red List; for example, *P. b. rabori* is considered Vulnerable^[21]^ and *P. b. iriomotensis* Critical Endangered^[22]^. The leopard cat in China is classified as Vulnerable^[23]^.

Most genetic studies of the leopard cat are from East Asia. The populations living on the islands of Japan have been classified into two subspecies: *P. b. euptilurus* and *P. b. iriomotensis*^[24-25]^. A study on population genetic structure revealed low levels of DRB allelic variation in MHC class II genes among subpopulations from the islands of Tsushima and Iriomote in Japan^[26]^, which is in line with findings of decreased genetic diversity based on DNA of the mitochondrial control region^[27]^.

Studies of leopard cats from Korea using microsatellite markers showed that this species had lower genetic diversity than 12 other feline species in the world^[28]^ and other mammalian species in Korea^[29]^. However, there has been little such research on species in China. One study found five management units for leopard cat populations based on genetic diversity and phylogenetic analysis using RAPD and variation in mitochondrial DNA^[30]^. Another study using mitochondrial DNA of Cyt B and the control region sequence found genetic diversity among populations from five geographic areas and detected independent trends in evolution between northern and southern branches in China^[31]^.

There is even less fieldwork on the status of wild mammals in the Beijing region. It is estimated that there were 1500 leopard cats in the first national wildlife survey from 1995 to 2000 (Report on resources survey on the terrestrial wild animals in Beijing, unpublished data, 2012). Other studies have detected food components of carnivores by scat residue identification^[32-33]^ and performed camera trapping monitoring and activity pattern analysis on wild mammals^[34-37]^. The only genetic study, on Chinese goral (*Naemorhedus griseus*) from the Songshan national nature reserve, confirmed 15 individuals and moderate population genetic diversity through noninvasive fecal sampling^[38]^. Clearly, there is a lack of research on the conservation of mammalian species in Beijing. Thus, we aimed to (1) clarify the genetic background of the leopard cat in Beijing through noninvasive fecal sampling and (2) detect genetic differentiation among subpopulations due to the segregation effects of infrastructure development and the expansion of human settlement. Given the high dispersal ability of the leopard cat and the fact that the influences of infrastructure development have only been at play for several decades, we assumed that there would be no genetic diversity among the subpopulations sampled.

## MATERIALS AND METHODS

### Sampling procedure and DNA extraction

From September 2017 to October 2018, we collected 601 fecal samples of suspected leopard cat origin by surveying transect lines from Songshan (n = 315), Yunmengshan (n = 43), Yunfengshan (n = 45), Xiaolongmen (n = 75), and Baihuashan (n = 123) nature reserves in Beijing (hereafter, SS, YMS, YFS, XLM, BHS, respectively; Fig 1). Fecal samples were put into sealed plastic bags, fixed with ethanol, and maintained at –20°C in the laboratory. We extracted total DNA from fecal samples using the Stool DNA Kit (D4015-01; Omega, Dorivalle, GA, USA) according to the manufacturer’s protocol.

**Figure 1.**
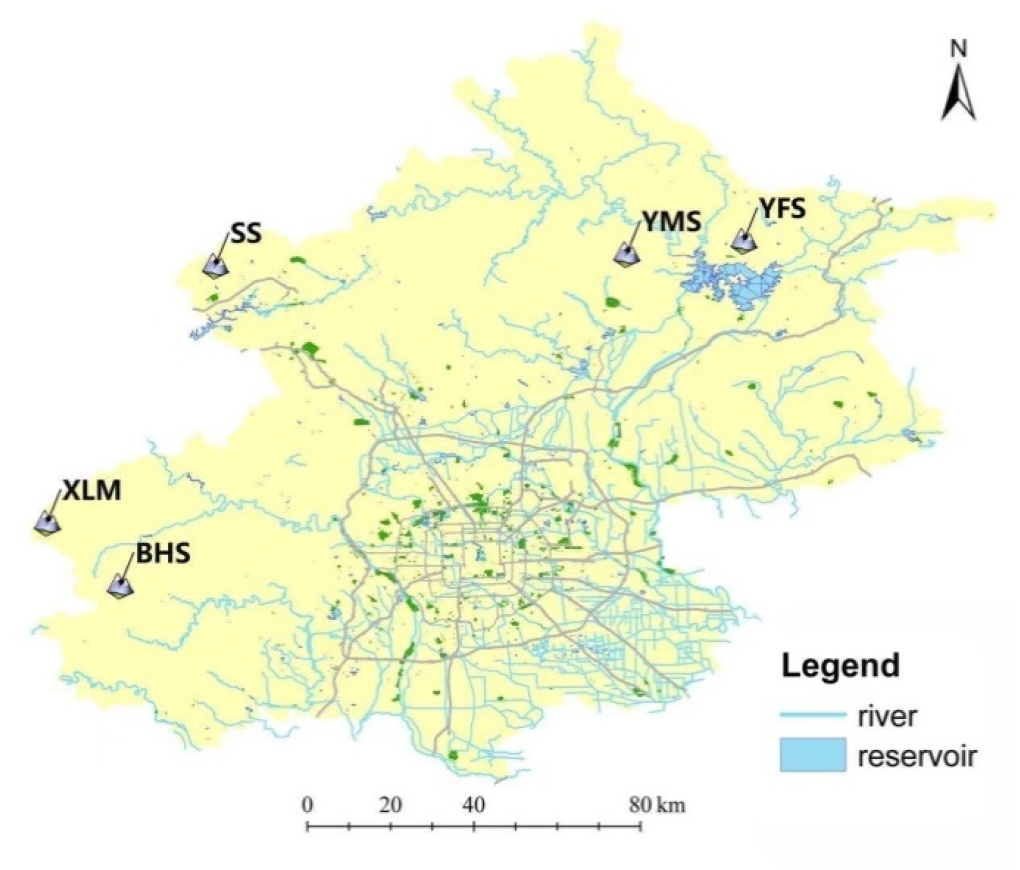
Fecal sampling areas for analyses of the genetic structure of leopard cats in Beijing BHS represents Baihuashan reserve, XLM represents Xiaolongmen reserve, SS represents Songshan reserve, YMS represents Yunmengshan reserve, and YFS represents Yunfengshan reserve.

### Species and sex identification

The universal primer of the carnivore species ATP6 hypervariable region of mitochondrial DNA was used in species identification^[39]^. PCR amplification was set to the following conditions: one cycle at 94°C for 5 min; 35 cycles at 94°C for 30 s, 60°C for 30 s, and 72°C for 45 s; and one cycle at 72°C for 8 min. Each 30 µL PCR reaction volume contained 15 µL Premix Ex Taq enzyme (Takara Biomedical Technology, Beijing, China), 0.2 µL bovine serum albumin (BSA), 1 µL forward or reverse primer, and 2 µL genomic DNA. The PCR products were purified before Sanger sequencing, and afterward one sequence was obtained (TsingKe Biotech, Beijing, China). The species was identified from a BLAST search of the NCBI database (GenBank no. KX857767.1) according to the degree of sequence matching. Sequences not matched to leopard cats were determined by the most closely matched source sequence. Each experiment included a positive and negative control (DNase/RNase-free deionized water template rather than DNA).

The zinc-finger regions of the X and Y chromosomes were used to identify the sex of the leopard cat fecal samples^[40]^. The PCR products of female samples were 165 bp, and the products of male samples were 162 and 165 bp. The 20 µL PCR reaction volume included 10 µL Premix Ex Taq enzyme (Takara Biomedical Technology), 0.2 µL BSA, 0.8 µL forward or reverse primer (the forward primer was labeled with FAM dye at the 5′ end), and 2 µL genomic DNA. The PCR conditions were 95°C for 10 min; 40 cycles at 95°C for 30 s, 58°C for 40 s, and 72°C for 30 s; and 72°C for 8 min. Genotyping of the samples was performed with an ABI 3730xl DNA analyzer (Applied Biosystems, Foster City, CA, USA) supplied by TsingKe Biotech. Each sample was analyzed at least three times until the exact genotype was obtained; samples without amplification products were excluded from the analysis.

### Microsatellite loci selection and amplification

We tried several reaction conditions and annealing temperatures repeatedly with fecal DNA samples based on 20 pairs of microsatellite loci for leopard cat from muscle samples^[28-29]^, and six suitable microsatellite loci (Pbe03, Pbe05, Pbe13, Pbe28, Pbe32, Pbe33) were selected. The conditions for the PCR reaction system were from Ko^[29]^. We genotyped the samples using the ABI 3730xl DNA analyzer supplied by TsingKe Biotech. Each sample was analyzed at least three times to reduce error. Samples amplified with six loci were considered to be successful and were used in the analysis.

### Amplification of the mitochondrial control region sequence

Two pairs of primers were designed to amplify the gene sequence of the mitochondrial control region of the identified leopard cat fecal DNA (Table 1) according to the mitochondrial genome sequence of the leopard cat in the NCBI database (GenBank no. KP246843.1). PCR amplification was as follows: one cycle at 94°C for 5 min; 35 cycles at 94°C for 30 s, 57°C or 60°C for 30 s, 72°C for 45 s; and one cycle at 72°C for 10 min. Each 30 µL PCR reaction volume contained 15 µL Premix Ex Taq enzyme (Takara Biomedical Technology), 0.2 µL BSA, 1 µL forward or reverse primer, and 2 µL genomic DNA. The PCR products were purified before Sanger sequencing, and afterward one sequence was obtained (TsingKe Biotech).

**Table 1.**
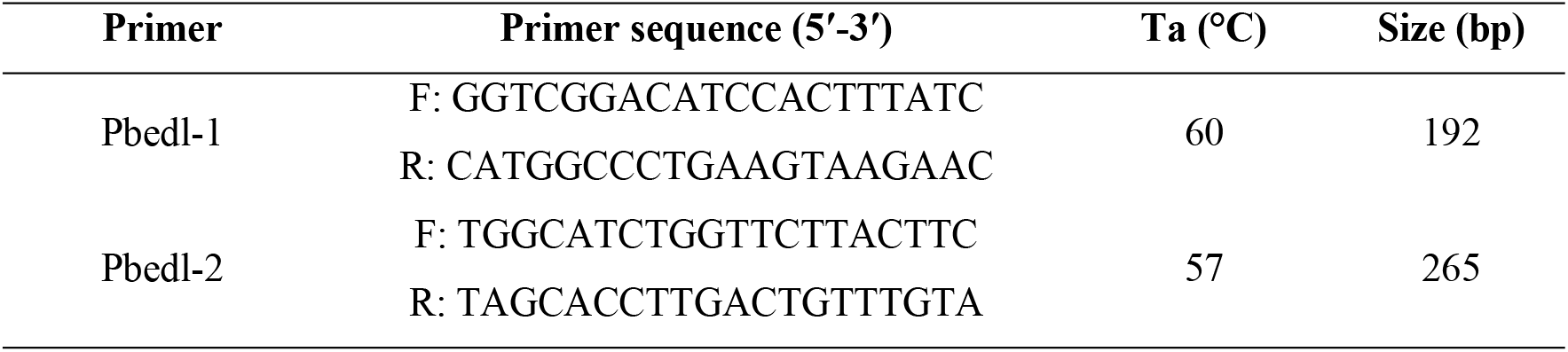
Primer pairs for amplifying the mitochondrial control region sequence of the leopard cat

### Data analysis

The sequenced DNA were checked with Clustal W v2.0 and manual proofreading, and ambiguous fragments and primer sequences were removed^[41]^. GeneMarker v2.2.0 was used to read the genotyping results^[42]^. We used Micro-Checker v2.2.3 to check for null alleles and genotyping errors^[43]^. The microsatellite data were organized in Microsoft Excel, and the MS tools plugin in Excel was used to find matching genotypes in the database. The reliability of the typing results was tested with the implementation criteria^[44]^. Samples were considered to originate from the same individual if the genotypes of all loci were identical or there was difference in only one locus^[45]^.

We tested the Hardy–Weinberg equilibrium and linkage disequilibrium of each target microsatellite locus using Arlequin v3.5^[46]^ and Genepop v4.3^[47]^ with Bonferroni correction. The number of alleles (Na), effective number of alleles (Ne), observed and expected heterozygosity (Ho and He, respectively), and polymorphism information content (PIC) were calculated with GenAlEx v6.5^[48]^. STRUCTURE v2.3.4 was used to verify and analyze the genetic differentiation among leopard cat subpopulations with Bayesian clustering^[49]^.

DnaSP v5.0^[50]^ was used to calculate the number of polymorphic loci (s), haplotype diversity (h), nucleotide diversity (π), and average number of different nucleotide (k) in the mitochondrial control region of the genome of the leopard cat population. The coefficient of genetic differentiation (Fst) among subpopulations was calculated with Arlequin v3.5.2.2^[46]^. Significance was tested with a random permutation test with 1000 replicates, and gene flow (Nm) was calculated based on Fst. MEGA v5 was used to calculate the genetic distance, base composition, gene sequence conversion, and transversion rate of leopard cats in the five sampling areas with Kimura’s two-parameter model^[51]^. The phylogenetic tree of the subpopulations in the five sampling areas was constructed by the neighbor-joining method with MEGA v5. The confidence of each branch of the tree was tested by 1000 bootstrap samples, and the mitochondrial control region sequence of the African lion (*Panthera leo*) in the NCBI database (KM374706.1) was used as the root of the phylogenetic tree. Tajima’s test was performed to determine whether the sequences conformed to neutral selection. NETWORK v5.0.0.1 was used to construct the haplotype network diagram for the leopard cat subpopulations by the median-joining method^[52]^.

## RESULTS

### Species and individual identification

Of the 601 samples collected in the five nature reserves, 550 were identified as coming from leopard cat, with matching rates ≥98% after a BLAST search of the NCBI database. Of those 550 samples, 508 were discriminated by sex, with 383 from females and 125 from males. The other samples were identified as red fox (*Vulpes vulpes*), Arctic fox (*Alopex lagopus*), domestic cat (*Felis catus*), and hog badger (*Arctonyx collaris*). The 508 samples successfully identified by sex were amplified with microsatellite loci; 96 samples that could not be amplified at the six sites were removed. Micro-Checker detected no invalid alleles at any of the six loci. After microsatellite genotype sharing analysis and sex identification, we identified 53 individuals (33 females and 20 males) in SS, 17 individuals (10 females and 7 males) in YMS, 17 individuals (14 females and 3 males) in YFS, 13 individuals (10 females and 3 males) in XLM, and 12 individuals (9 females and 3 males) in BHS.

### Microsatellite analysis of genetic diversity

None of the six microsatellite loci used for the analysis of genetic diversity deviated from Hardy-Weinberg equilibrium (P > 0.05), and no linkage disequilibrium was detected. The average Na of the six microsatellite loci was 3.400, the average Ne was 2.288, the average Ho was 0.392, the average He was 0.514, and the average PIC was 0.449 (Table 2).

**Table 2.** Genetic diversity of six microsatellite loci detected in leopard cat subpopulations in the five sampling areas (average)

| Microsatellite<br>index | SS<br>(N = 53) | YMS<br>(N = 17) | YFS<br>(N = 17) | XLM<br>(N = 13) | BHS<br>(N = 12) |
| --- | --- | --- | --- | --- | --- |
| Na | 4.333 | 2.667 | 3.500 | 3.167 | 3.333 |
| Ne | 2.580 | 2.087 | 2.354 | 2.271 | 2.147 |
| Ho | 0.531 | 0.353 | 0.392 | 0.295 | 0.389 |
| He | 0.535 | 0.498 | 0.517 | 0.534 | 0.484 |
| PIC | 0.470 | 0.418 | 0.452 | 0.470 | 0.434 |

The Fst of the leopard cat subpopulations in five sampling areas ranged from 0.011 to 0.082; Fst values of the subpopulations in BHS and YMS, BHS and YFS, SS and YMS, and SS and BHS ranged from 0.053 to 0.082, showing little genetic differentiation among these subpopulations (Table 3). Meanwhile, Nm was relatively large, ranging from 2.799 to 22.478, showing normal gene flow among subpopulations.

**Table 3.** Fst and Nm based on analyses of the diversity of microsatellite loci in leopard cat subpopulations in the five sampling areas

| Sampling area | SS | YMS | YFS | XLM | BHS |
| --- | --- | --- | --- | --- | --- |
| SS |  | 4.136 | 18.981 | 11.655 | 2.799 |
| YMS | 0.057 |  | 4.852 | 9.366 | 4.467 |
| YFS | 0.013 | 0.049 |  | 22.478 | 3.222 |
| XLM | 0.021 | 0.026 | 0.011 |  | 6.695 |
| BHS | 0.082 | 0.053 | 0.072 | 0.036 |  |

Fst appears below the diagonal, and Nm appears above the diagonal.
SS represents Songshan reserve, YMS represents Yunmengshan reserve, YFS represents Yunfengshan reserve, XLM represents Xiaolongmen reserve, and BHS represents Baihuashan reserve. The same in the following tables.

The results of STRUCTURE analyses showed that when k = 1, k = 2, k = 3, k = 4, and k = 5, there was no significant genetic differentiation among the subpopulations in the five sampling areas. This indicates that the leopard cats maintained gene exchange among different distribution ranges (Fig 2).

**Figure 2.**
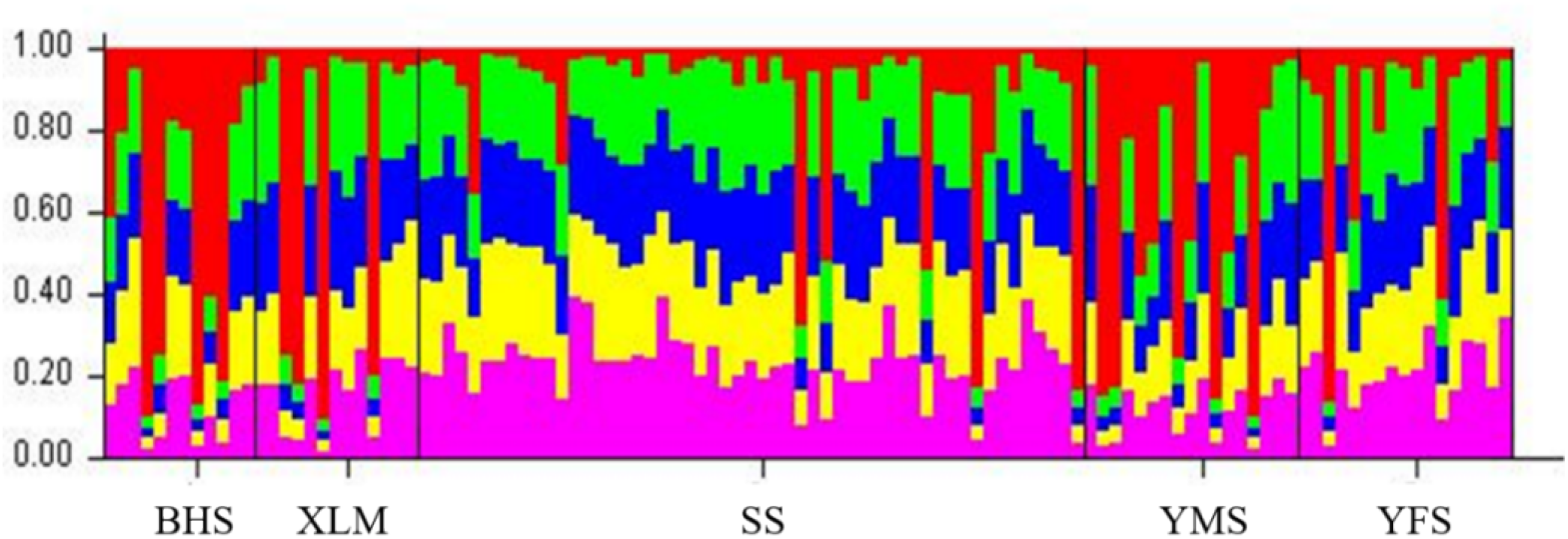
Genetic structure based on microsatellite genotypes in leopard cat subpopulations in the five sampling areas (k = 5) BHS represents Baihuashan in orange, XLM represents Xiaolongmen in dark blue, SS represents Songshan in green, YMS represents Yunmengshan in dark green, and YFS represents Yunfengshan in purple.

### Analysis of the genetic diversity of the mitochondrial DNA control region

After individual identification by microsatellite genotyping, confirmed samples were amplified with the mitochondrial control region primer and 550 bp fragments of the partial sequences were obtained. All sequences were tested with Tajima’s test, and the difference was not significant (P > 0.10), which conforming to neutral selection.

Among the genetic diversity parameters of the mitochondrial control region calculated by DnaSP v5.0, the n of the leopard cat subpopulations in the five sampling areas ranged from 7 to 34, s ranged from 18 to 61, and h was higher at SS and BHS (Table 4). π was higher at YMS and YFS and was larger than 0.01. k ranged from 5.505 to 13.020.

**Table 4.** Diversity of the mitochondrial DNA control region in leopard cat subpopulations in the five sampling areas

| Sampling area | $n$ | $s$ | $h$ | $\pi$ | $k$ |
| --- | --- | --- | --- | --- | --- |
| SS | 34 | 40 | $0.978 \pm 0.012$ | $0.005686 \pm 0.000329$ | 13.020 |
| YMS | 6 | 61 | $0.675 \pm 0.117$ | $0.013850 \pm 0.006886$ | 10.942 |
| YFS | 8 | 18 | $0.838 \pm 0.085$ | $0.015728 \pm 0.004945$ | 5.505 |
| XLM | 7 | 35 | $0.731 \pm 0.133$ | $0.005870 \pm 0.002726$ | 6.692 |
| BHS | 10 | 25 | $0.970 \pm 0.014$ | $0.005195 \pm 0.001038$ | 6.909 |

According to the genetic diversity of the mitochondrial control region sequences, the genetic distance between the BHS and XLM subpopulations was 0, which indicated that these two subpopulations could be included in one group, and the genetic distance between the SS and YFS subpopulations was the largest at 0.3434 (Table 5).

**Table 5.** Genetic distance between leopard cat subpopulations in the five sampling areas based on the mitochondrial DNA control region sequence

| Sampling area | SS | YMS | YFS | XLM |
| --- | --- | --- | --- | --- |
| YMS | 0.2783 |  |  |  |
| YFS | 0.3434 | 0.0866 |  |  |
| XLM | 0.2296 | 0.0415 | 0.0495 |  |
| BHS | 0.2296 | 0.0415 | 0.0495 | 0 |

Fst between YMS and BHS, YMS and XLM, and BHS and XLM was 0; thus, the gene flow (Nm) among these areas was nonsignificant. The BHS and XLM subpopulations had the same Fst and Nm values as those in the other three sampling areas, which shows that there was no genetic differentiation in these two areas. Fst between SS and YMS and between YFS and BHS/XLM was larger than 0.05, showing a trend toward genetic differentiation (Table 6).

**Table 6.**
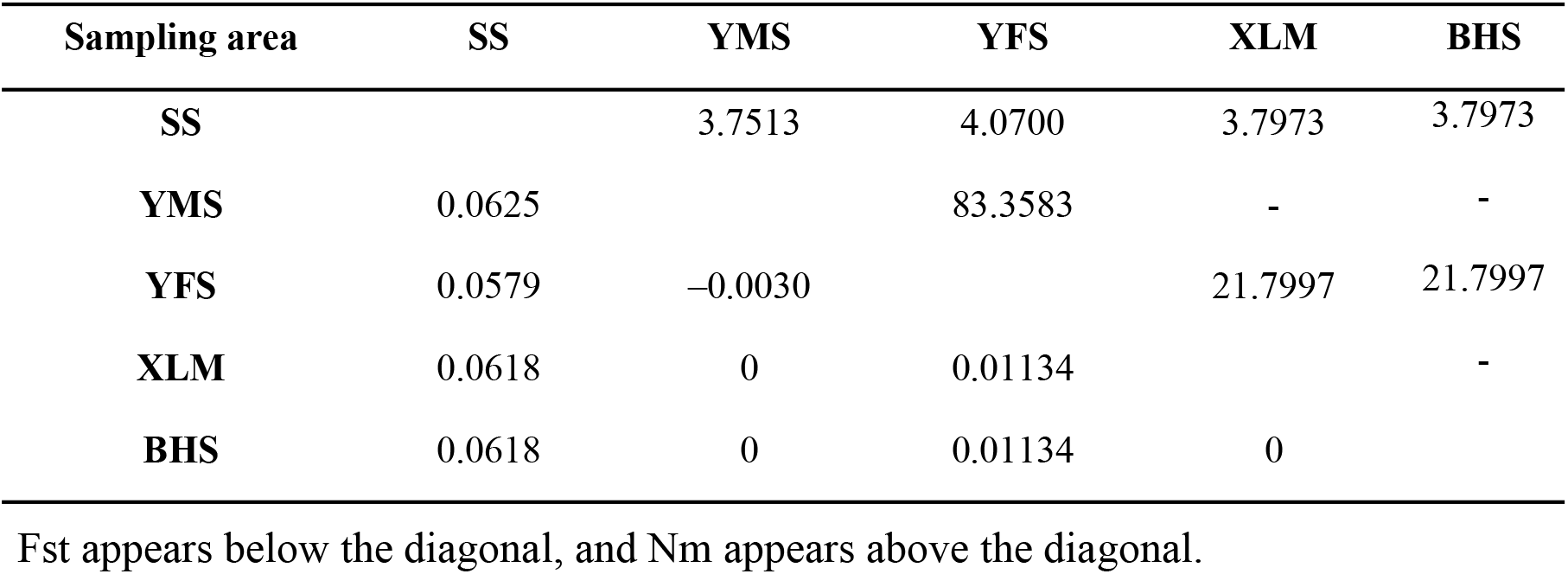
Fst and Nm in leopard cat subpopulations in the five sampling areas based on the mitochondrial DNA control region

| Sampling area | SS | YMS | YFS | XLM | BHS |
| --- | --- | --- | --- | --- | --- |
| SS |  | 3.7513 | 4.0700 | 3.7973 | 3.7973 |
| YMS | 0.0625 |  | 83.3583 | - | - |
| YFS | 0.0579 | -0.0030 |  | 21.7997 | 21.7997 |
| XLM | 0.0618 | 0 | 0.01134 |  | - |
| BHS | 0.0618 | 0 | 0.01134 | 0 |  |

Fst appears below the diagonal, and Nm appears above the diagonal.

A total of 11 haplotypes were defined from the 101 partial sequences of the mitochondrial control regions amplified from the five sampling areas (GenBank access nos. MW648825–MW648926). The network diagram of haplotypes among the five subpopulations showed that most came from the SS subpopulation. One of these haplotypes (Hap_8) was shared by the five subpopulations and had the highest frequency; Hap_1 was shared by three subpopulations (SS, YFS, and YMS; Fig 3).

**Figure 3.**
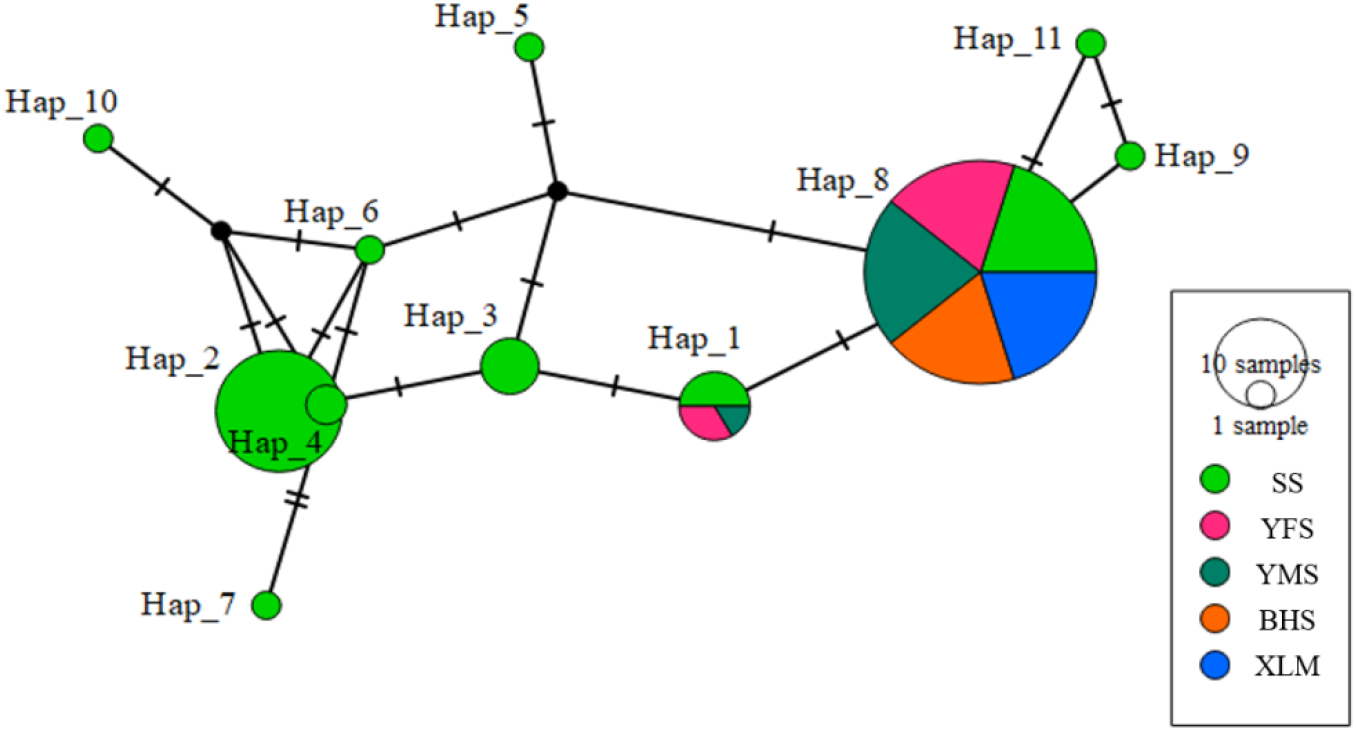
Haplotype network based on mitochondrial DNA control region in leopard cat subpopulations in the five sampling areas The different patterns in the circles show the distribution of haplotypes in different subpopulations, the size of the circle represents the haplotype frequency, and the hash marks indicate mutation between haplotypes.

Based on phylogenetic tree analysis of the partial sequences of the mitochondrial control regions, the five subpopulations were divided into two branches: 28 individuals all from SS made up one branch, and the other four subpopulations and the remaining SS individuals were clustered into another branch (Fig 4). This is consistent with the results of Fst and Nm analyses and implies that the SS subpopulation tended to differentiate from the other groups.

**Figure 4.**
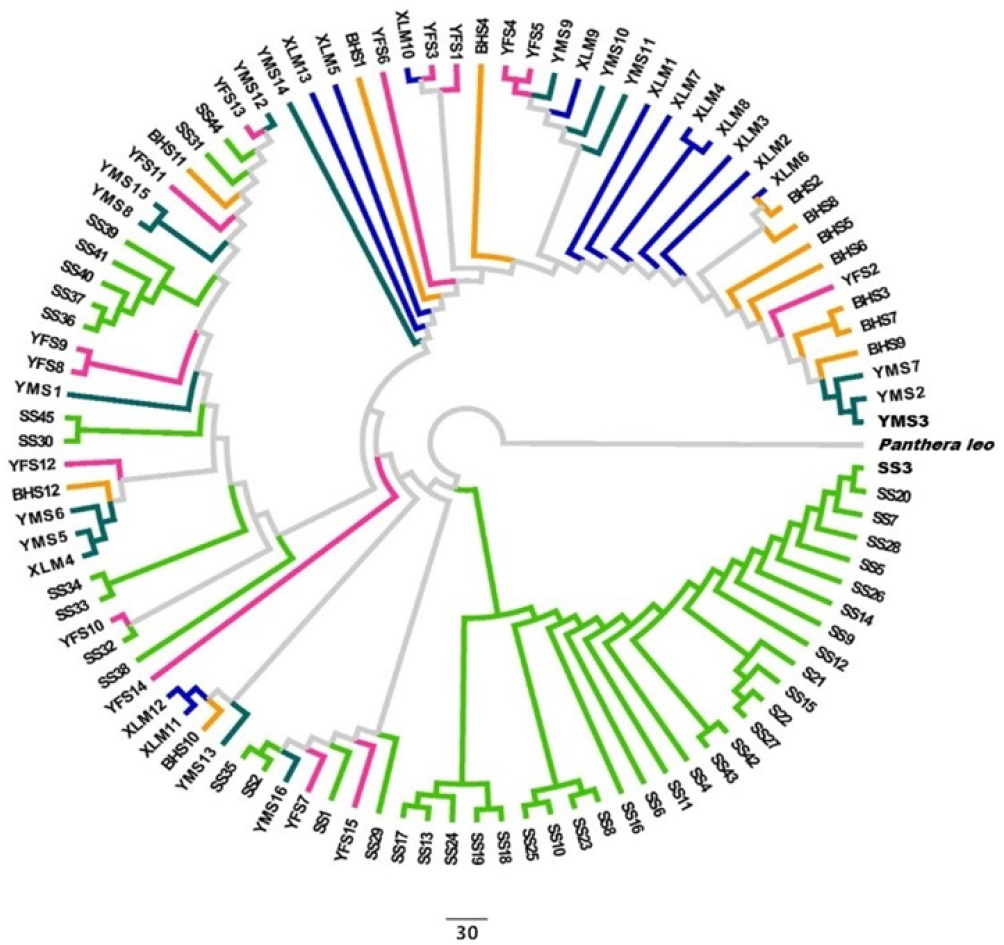
Phylogenetic tree of leopard cat subpopulations based on partial sequences of the mitochondrial DNA control region in the five sampling areas SS represents Songshan in green, YMS represents Yunmengshan in dark green, YFS represents Yunfengshan in purple, XLM represents Xiaolongmen in dark blue, and BHS represents Baihuashan in orange. *Panthera leo* is the outgroup.

## DISCUSSION

Accurate evaluation of population abundance and sex structure is critical to assessing the current status and future development of local populations^[53]^. From large samples of scat, we identified 112 leopard cats from five nature reserves in mountainous habitats around Beijing. Most individuals were from the SS reserve. The 53 leopard cats in the 45 km^2^ SS reserve^[54]^, may approximate the real state of the population, whereas the abundance in the other reserves may be underestimated because of the lack of scat samples. We suggest further field sampling to obtain a clearer estimation of the leopard cat population in the Beijing region.

The sex ratio is an important indicator of population structure and development trends for effective management of endangered species^[55]^. There was a general trend toward a female-biased sex ratio in the five sampling areas (the female-to-male ratio was 1.65:1 at SS, 1.43:1 at YMS, 4.67:1 at YFS, 3.33:1 at XLM, and 3:1 at BHS). This is in line with the female-biased sex structures of other feline species, such as the jungle cat (*Felis chaus*) in India at 1.59:1^[56]^. This phenomenon is more obvious among large cats such as the snow leopard (*Panthera uncia*) and African lion (*Panthera leo*;) ^[57-58]^ because males die at higher rates than females from injuries from hunting and individual competition^[57,59]^. However, this female-biased sex ratio may help the population recover quickly from low numbers^[60]^. Our study provides valuable reference information for future monitoring of the leopard cat sex ratio in the Beijing region.

### Genetic structure based on microsatellite diversity

Analyzing the genetic structure of local populations through molecular approaches may reveal clues to their evolutionary status and adaptability to the environment^[61]^. Microsatellites are frequently selected as nuclear markers for their high polymorphism, low requirements for DNA in PCR amplification, and accuracy in assessing population genetic diversity^[28]^. In our study the average PIC at all six microsatellite loci was above 0.4 (Table 2), which indicates moderate diversification of alleles according to the standards of Botstein^[62]^. In this study, the average effective number of alleles (Ne = 2.29) was lower than the index for leopard cat in Korea (Na = 3.8) ^[29]^ but higher than that of the Iriomote leopard cat population in Japan (Na = 1.33) ^[26]^. Moreover, the average observed heterozygosity was lower than in Korea (Ho = He = 0.41) ^[29]^, but the average expected heterozygosity was higher (Table 2), and compared to the Tsushima leopard cat population in Japan (Ho = 0.77, He = 0.66)^[26]^ our parameters were much lower. Thus, total genetic diversity was moderate in this study. We postulate the existence of population shrinkage from overhunting^[63]^ and thereafter population expansion for the leopard cat population in the Beijing region.

### Genetic structure based on diversity of the mitochondrial DNA control region

The evolutionary rate of control region is the fastest of mitochondrial DNA genome, and its sequence diversity is frequently used in studies of population genetics^[64]^. We detected 11 haplotypes based on 550 bp partial sequences of the control region, of which 9 were typical of the SS subpopulation and only one was shared by the five subpopulations (Fig 3). This may imply a discrepancy between the SS subpopulation and the others, or it may be that more cat individuals were identified in this sampling area than in the others. Further work is needed to clarify this.

Genetic diversity is the fundamental factor determining species adaptability and one of the core topics in biodiversity research. In this study the h of mitochondrial DNA ranged from 0.675 to 0.978 and π ranged from 0.005195 to 0.015728 (Table 4), in the moderate range for leopard cat populations in Asia (h = 0.834 ± 0.035, π = 0.0075 ± 0.0004) ^[65]^. Of the five subpopulations, YMS and YFS had higher nucleotide diversity than the others, which may indicate that the two groups were in good genetic condition. There were large differences among the five subpopulations in terms of haplotype diversity and nucleotide diversity. We assume that this is related to differential effects of human disturbance in terms of time and magnitude, such as the building of tourism facilities, road construction, and the expansion of human settlement.

### Population genetic differentiation

Habitat loss and fragmentation increase the chance for inbreeding and pose a risk for extinction on local populations, which is becoming a major threat to biodiversity conservation^[58]^. In general, wild animals must have a high rate of dispersal to reduce this risk and maintain a diversified population genetic structure^[14]^. Based on an analysis of the diversity of microsatellite loci, we found frequent gene flow among the five leopard cat subpopulations, with Nm between 5.598 and 44.955. The STRUCTURE diagram also revealed gene exchange in the five sampling areas, indicating no genetic discrepancy (Fig 2). However, the results of gene flow analysis with partial sequences of the mitochondrial DNA control region demonstrated that Fst was between 0.05 and 0.15 in SS and YMS, SS and BHS, YMS and BHS, and YFS and BHS, respectively, revealing moderate differentiation among these subpopulations (Table 6).

At the same time, the phylogenetic tree exhibited two branches for the five subpopulations, with SS individuals making up one branch (Fig 4), which may imply that the SS subpopulation has diverged from the other groups. Thus, greater attention must be paid to this subpopulation in future monitoring programs, especially because the 2022 Winter Olympic Games will be held nearby and the influence of the construction of game facilities on individual dispersion, habitat occupation, and genetic structure must be considered.

## CONCLUSION

From the point of view of conservation genetics, the Beijing region is in a relatively narrow geographic range with no marked differences in weather or altitude. In addition, the leopard cat has strong dispersal ability. We postulated at the beginning of our study that there would be no genetic diversity among populations of leopard cat in this area. However, the results indicated that the SS subpopulation was different from the others. We assume that this is due to segregation effects from natural rivers, major roads, and the expansion of human residential sites. Thus, we suggest that the SS subpopulation be taken as an independent conservation unit for the planning of further conservation strategies. If needed, individuals could be introduced to maintain the genetic diversity of the local population.

## AUTHOR CONTRIBUTIONS

L.X.Gai, J.Li, and W.D.Bao, designed the experiments; Y.Teng, and J.Yang, performed the experiments and wrote the paper; F.L.G, and L.F.Ju, analyzed the data. All authors read and approved the final version of the manuscript.

## ACKNOWLEDGMENTS

We thank the staff of the nature reserves for help collecting the fecal samples.

